# The SUMO ligase Su(var)2-10 controls eu- and heterochromatic gene expression via establishment of H3K9 trimethylation and negative feedback regulation

**DOI:** 10.1101/533232

**Authors:** Maria Ninova, Baira Godneeva, Yung-Chia Ariel Chen, Yicheng Luo, Sharan J. Prakash, Ferenc Jankovics, Miklós Erdélyi, Katalin Fejes Tóth, Alexei A. Aravin

**Affiliations:** California Institute of Technology, Division of Biology and Biological Engineering, 147-75 1200 E. California Blvd., Pasadena, CA 91125, USA; Institute of Molecular Genetics, Russian Academy of Sciences, Moscow 123182, Russia; institute of Genetics, Biological Research Centre of the Hungarian Academy of Sciences, Szeged 6726, Hungary

**Keywords:** chromatin, heterochromatin, epigenetics, gene regulation, transcriptional repression, transposons, germline, cell fate maitanance

## Abstract

Chromatin is critical for genome compaction and gene expression. On a coarse scale, the genome is divided into euchromatin, which harbors the majority of genes and is enriched in active chromatin marks, and heterochromatin, which is gene-poor but repeat-rich. The conserved molecular hallmark of heterochromatin is the H3K9me3 modification, which is associated with gene silencing. We found that in *Drosophila* deposition of most of the H3K9me3 mark depends on SUMO and the SUMO-ligase Su(var)2-10, which recruits the histone methyltransferase complex SetDB1/Wde. In addition to repressing repeats, H3K9me3 also influences expression of both hetero- and euchromatic host genes. High H3K9me3 levels in heterochromatin are required to suppress spurious non-canonical transcription and ensure proper gene expression. In euchromatin, a set of conserved genes is repressed by Su(var)2-10/SetDB1-induced H3K9 trimethylation ensuring tissue-specific gene expression. Several components of heterochromatin are themselves repressed by this pathway providing a negative feedback mechanism to ensure chromatin homeostasis.

**Highlights:**

- Proper expression of host genes residing in heterochromatin requires Su(var)2-10-dependent installation of the H3K9me3 mark to suppress spurious non-canonical transcription.
- A set of euchromatic host genes is repressed by transposon-independent installation of H3K9me3 in a process that depends on Su(var)2-10 and SUMO.
- Installation of H3K9me3 via Su(var)2-10 ensures tissue-specific gene expression.
- H3K9me3-dependent silencing of genes encoding proteins involved in heterochromatin formation provides negative feedback regulation to maintain heterochromatin homeostasis.

## Introduction

Eukaryotic DNA is packaged in nucleosomes comprised of DNA and histone octamers to form chromatin. Post-translational modifications of histone proteins regulate chromatin compaction, which is critical for nuclear architecture and essential processes such gene expression control, recombination, replication, and DNA repair (reviewed in Bannister and Kouzarides, 2011). Chromatin has historically been differentiated into euchromatin and heterochromatin based on density reflected by differential staining (Heitz, 1928). Euchromatin is relatively uncondensed, ‘open’ chromatin where transcription is active, whereas heterochromatin is compact, and transcription is repressed in these regions of DNA. Heterochromatin regions are relatively gene-poor and repeat-rich, including satellite repeats concentrated around chromosomal arm ends, as well as transposable elements (TEs) – parasitic DNA elements capable of copying/cutting and integrating their sequences within host genomes (Allshire and Madhani, 2018; Henikoff, 2000). Such loci are typically condensed in all cell types and are referred to as ‘constitutive heterochromatin’. Constitutive heterochromatin is essential for genome stability, as it prevents illicit recombination between repetitive regions (Janssen et al., 2018; Peng and Karpen, 2009). ‘Facultative’ heterochromatin includes regions repressed in a cell-type specific manner, such as genes that are switched off during differentiation. Of note, although considered transcriptionally silenced, certain regions of heterochromatin, such as rRNA loci, are highly transcribed (Gatti and Pimpinelli, 1992; Weiler and Wakimoto, 1995; Yasuhara and Wakimoto, 2006). In *Drosophila*, some genes residing in heterochromatin in fact require this environment to be properly expressed (Eberl et al., 1993; Lu et al., 1996, 2000; Wakimoto et al., 1990; Yasuhara and Wakimoto, 2006), although the molecular mechanism of this phenomenon remains elusive.

In the past several decades, significant efforts have been dedicated to identifying factors that regulate the deposition, erasure, and recognition of different histone modifications and to elucidating the biological functions of such marks. The best studied repressive marks associated with heterochromatin are histone 3 lysine 9 trimethylation (H3K9me3) and H3 lysine 27 trimethylation (H3K27me3). H3K27me3 is deposited by the polycomb repressive complex 2 (PRC2) and is primarily involved in facultative heterochromatin formation (Margueron and Reinberg, 2011; Schuettengruber et al., 2017). PRC2/H3K27me3 silencing targets a large numbers of developmental genes such as the homeobox complex genes (Margueron and Reinberg, 2011; Schuettengruber et al., 2017). H3K9me3 is a conserved hallmark of constitutive heterochromatin, highly enriched at centrometric and telomeric repeats from yeast to human, and has a well-established role in transposon silencing in metazoans (Karimi et al., 2011; LeThomas et al., 2013; Martens et al., 2005; Matsui et al., 2010; Mikkelsen et al., 2007; Pezic et al., 2014; Rozhkov et al., 2013; Sienski et al., 2012). H3K9me3 is deposited by a conserved class of SET-domain containing K9-specific histone methyltransferase enzymes (HMTs), such as SETDB1 and SUV39 (Nakayama, 2001; Rea et al., 2000; Schultz et al., 2002), and provides a high-affinity binding site for members of the conserved heterochromatin protein 1 (HP1) family (Bannister et al., 2001; Jacobs et al., 2001; Lachner et al., 2001). HP1 proteins can oligomerize, resulting in local chromatin compaction, and can further recruit additional H3K9me3 writer complexes enabling heterochromatin spreading (Canzio et al., 2011; Hiragami-Hamada et al., 2016). Thus, H3K9me3 deposition is a critical step in heterochromatin regulation.

Efforts to understand the recruitment of H3K9 methyltransferases to target loci have revealed a number of different pathways that employ a diversity of guides such as DNA binding proteins and non-coding RNAs. For example, in yeast small interfering RNAs (siRNAs) and Argonaute proteins are required for recruitment of the silencing machinery to pericentric heterochromatin, while DNA-binding factors that recognize specific regulatory elements can also nucleate heterochromatin formation at the mating type locus and at telomeres (Hall, 2002; Jia, 2004; Kanoh et al., 2005; Verdel et al., 2004; Volpe et al., 2002). In the germ cells and ovarian somatic cells of *Drosophila*, Piwi-associated RNAs (piRNAs), a dedicated class of small RNAs antisense to TEs, direct H3K9 trimethylation at TE loci (LeThomas et al., 2013; Rozhkov et al., 2013; Sienski et al., 2012). Similarly, piRNAs and Piwi proteins are required for H3K9me3 deposition and LINE element silencing in mouse germ cells (Pezic et al., 2014). The vertebrate-specific Krüppel-associated box-containing zinc finger proteins (KRAB-ZFPs) are another class of HMT guides. KRAB-ZFPs recognize specific DNA motifs of endogenous retrovirus and recruit the mammalian homolog of dSetDB1, ESET/SETDB1, resulting in H3K9me3 deposition and silencing of endogenous retrovirus regions (Wolf et al., 2015). Thus, heterochromatin can be established by different molecular pathways in different organisms, different cell types, and even at different genomic loci within the same cell. A complete picture of the principles and molecular mechanisms that recruit H3K9me3 methyltransferases to different target loci is yet to be established.

We have identified the conserved SUMO E3 ligase Su(var)2-10/dPIAS as a novel factor essential for H3K9me3 deposition in *Drosophila.* The Su(var)2-10 locus was originally identified in a genetic screen as a suppressor of positional effect variegation – a phenomenon in which euchromatic genes translocated near heterochromatin are silenced (Elgin and Reuter, 2013; Reuter and Wolff, 1981), strongly suggesting that it is important in heterochromatin formation. Further, Su(var)2-10 is required for chromosomal stability (Hari et al., 2001). In the accompanying manuscript, we report that Su(var)2-10 is required for genome-wide H3K9me3 deposition and transcriptional repression of TEs in female germ cells. In female germ cells, Su(var)2-10 interacts with and acts downstream of the piRNA-Piwi complex, and its SUMO ligase activity is required to recruit the SetDB1 silencing complex to target TE loci (Ninova et al., accompanying manuscript). Reduction in Su(var)2-10 levels in germ cells affected the vast majority of H3K9me3-rich domains, and also affected the expression of multiple host genes. Here we describe the distinct mechanisms though which H3K9me3 deposition is involved in gene regulation. First, we found that host gene expression can be altered as a result of H3K9me3 spreading from adjacent TEs, demonstrating that novel TE integrations are an important source of epigenetic diversity. Second, we showed that the H3K9me3 mark and Su(var)2-10 are required for the activity of genes residing in heterochromatin, likely by preventing interfering transcription from adjacent TEs. Furthermore, H3K9me3 is required to repress cryptic promoters upstream or within host gene introns, thereby suppressing aberrant gene isoforms. Finally, we showed that Su(var)2-10/SUMO-dependent H3K9me3 islands regulate a subset of host genes without proximal TE insertions, indicating the existence of piRNA-independent H3K9me3 deposition. Targets of piRNA-independent H3K9me3 deposition include protein-coding genes and non-coding RNAs normally expressed in other tissues, highlighting the role of this mark in cell lineage commitment. Remarkably, several genes that encode proteins required for heterochromatin formation are also the targets of heterochromatin repression pointing to a negative feedback mechanism of heterochromatin control.

## Results

### Su(var)2-10 controls the expression of host genes

We have shown that Su(var)2-10 represses transposons recognized by the Piwi/piRNA complex (Ninova et al., accompanying manuscript). However, in contrast to Piwi and piRNA, which are predominantly expressed in the germline and required for fertility but not somatic development, Su(var)2-10 is expressed ubiquitously and null-mutants are lethal, suggesting that Su(var)2-10 might have other, piRNA-independent, targets and functions. To identify genes that are regulated by Su(var)2-10, we inhibited expression of Su(var)2-10 in ovarian germ cells (GLKD) using small hairpin RNA (shRNA) driven by the maternal tubulin GAL4 driver (mtGAL4) and analyzed changes in steady state RNA levels in shSu(var)2-10-expressing and control shWhite lines (shW). Differential gene expression analysis of RNA-seq libraries performed in duplicates showed that 662 and 417 genes were more than two-fold up- and down-regulated, respectively, upon depletion of Su(var)2-10 (FDR 5%, DESeq2; Fig. 1A). Thus, Su(var)2-10 regulates the expression of a subset of host genes in addition to its function in transposon silencing described in the accompanying manuscript (Ninova et al., accompanying manuscript).

**Figure 1.**
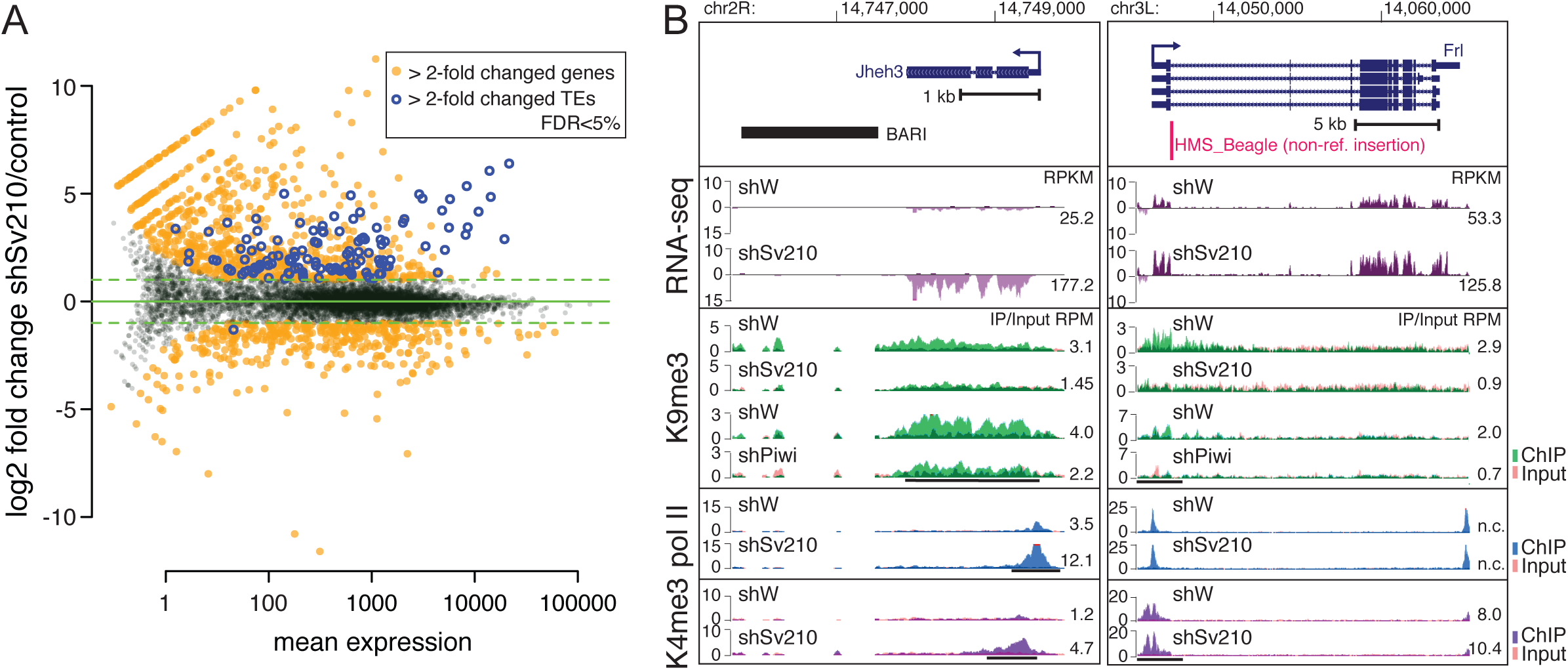
Su(var)2-10 depletion leads to transcriptome-wide changes in gene expression that correlate with changes in chromatin marks. (A) Differential gene expression analysis of Su(var)2-10 GLKD vs. control (shW) ovaries. Scatter plot of the mean expression of genes and transposons (x-axis) and log2-normalized fold change between Su(var)2-10 KD and control (y-axis). Significantly up- or down-regulated genes and TEs (>2-fold change, FDR<5%; DESeq2) are highlighted. Data are from two biological replicates. (B) UCSC browser tracks show examples of euchromatic genes adjacent to reference (left) and non-reference (right) TE insertions sensitive to Su(var)2-10 GLKD. Histograms of the normalized signal from RNA-seq and ChIP-seq data, as indicated, from Su(var)2-10 KD and control ovaries. For ChIP-seq tracks, ChIP and Input signals are overlaid. Top panels show the gene structures at corresponding loci. The position of the nonreference insertion is marked by a red line. Arrows indicate the direction of transcription. Numbers on the right are the RPKM values of exonic regions (RNA-seq, estimated by eXpress) or normalized ChIP/Input signal (ChIP-seq) in manually selected genomic intervals indicated by black bars.

### Spreading of the H3K9me3 mark from transposon sequences leads to repression of adjacent genes

We showed that silencing of transposons by Su(var)2-10 involves deposition of the H3K9me3 mark by the SetDB1/Wde histone methyltransferase complex (Ninova et al., accompanying manuscript). Repressive chromatin marks can spread from transposon sequences up to several kilobases (kb) and influence the expression of adjacent host genes (Lee and Karpen, 2017; Pezic et al., 2014; Sentmanat and Elgin, 2012; Sienski et al., 2012). We found that several genes regulated by Su(var)2-10 harbor or are adjacent to transposon insertions present in the reference genome sequence. For example, we observed H3K9me3 spreading from the *BARI* element at the *jheh* locus into the downstream *jheh3* gene. GLKD of Su(var)2-10 led to reduced H3K9me3 levels and increased expression of *jheh3* in the ovary (Fig. 1B, left). This result indicates that the transposon insertion influences *jheh3* expression through its effect on the local chromatin state rather than through disruption of gene regulatory elements in the DNA sequence. The *BARI* insertion at the *jheh3* locus was shown to be positively selected in the *D. melanogaster* population (Gonzalez et al., 2009; González et al., 2008) indicating that Su(var)2-10-dependent repression caused by a transposon insertion can be used for beneficial re-wiring of host gene regulatory networks.

Many genes regulated by Su(var)2-10 do not have transposon insertions in their vicinities in the reference genome sequence. However, previous studies that mapped TE insertions in various *D. melanogaster* strains revealed that many TE insertions are polymorphic, i.e., they are present in some strains and absent in others (Rahman et al., 2015). As the strain used in our study is different from the strain used for the reference genome assembly, we mapped non-reference transposon insertions in our strain using the TIDAL pipeline (Rahman et al., 2015). This analysis revealed over 400 non-reference TE insertions common to our control and to the Su(var)2-10 GLKD strain. Similar to transposon sequences present in the reference genome, many non-reference TE insertions are targets of piRNA-dependent silencing and show H3K9me3 enrichment that is dependent on Su(var)2-10 (Ninova et al., accompanying manuscript). We found that several host genes are affected by Su(var)2-10 knockdown as a consequence of H3K9me3 deposition on non-reference TEs; an example is the *Frl/CG43986* gene (Fig. 1B, right). Thus, spreading of repressive chromatin from both reference and non-reference TE insertions can lead to repression of adjacent host genes confirming that transposons contribute to regulation of host gene expression. Importantly, our results show that transposons do not need to disrupt cis-regulatory DNA sequences in order to influence gene expression. Instead, spreading of chromatin marks from a transposon into gene regulatory elements can result in repression.

### Su(var)2-10-dependent H3K9me3 installation ensures proper expression levels and isoform selection of heterochromatic genes

Though new transposon insertions often occur in gene-dense regions in euchromatin, the vast majority of transposons accumulate in heterochromatin. The heterochromatic compartment of *D. melanogaster* is well characterized cytologically and through genome-wide profiling of chromatin marks and includes nearly the entire chromosomes Y and 4 as well as pericentromeric and telomeric regions of chromosomes X, 2, and 3 (Gatti and Pimpinelli, 1992; Hoskins et al., 2002, 2007; Riddle et al., 2011). Heterochromatin-euchromatin boundaries are similar in different tissues and developmental stages (Riddle et al., 2011; see methods for heterochromatin domain annotation), and our H3K9me3 ChIP-seq profiles from ovaries of control flies (shW) largely overlap with annotated domains (Fig. 2A). Despite its relatively low gene density, several hundred protein-coding genes reside in heterochromatin (Hoskins et al., 2002): Of the 8255 protein-coding genes expressed in the ovary, 228 reside in heterochromatin including genes that encode conserved proteins with well-established functions such as *Parp* encoding poly-(ADP-ribose) polymerase and *AGO3* required for piRNA repression.

**Figure 2.**
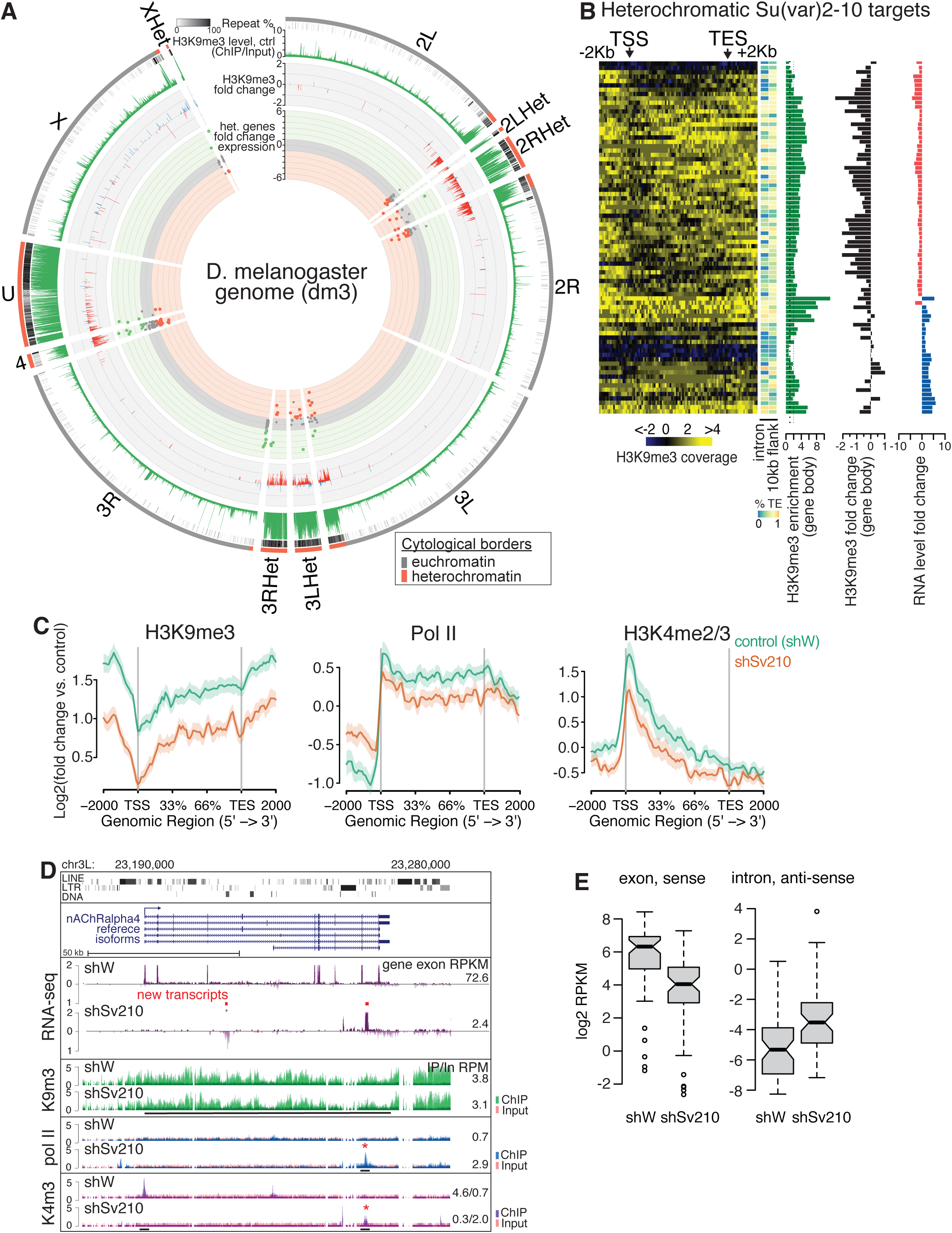
Su(var)2-10 and H3K9me3 are required for heterochromatic gene expression. (A) Epigenetic landscape of heterochromatic Su(var)2-10 target genes. Circle plot of the *D. melanogaster* chromosome arms and heterochromatic scaffolds (dm3 assembly). Tracks show, from outer to inner circles, orientations of chromosomal arms and scaffolds with annotated heterochromatic regions (red, see Methods), repetitive element percentage at 5-kb genomic intervals, H3K9me3 signal in control (shW) ovaries calculated for 5-kb genomic windows as log2-normalized ratio of ChIP-seq to Input RPKM values, log2-transformed fold change of H3K9me3 signal in Su(var)2-10 GLKD versus control (shW) ovaries for regions with >2 fold enrichment in control ovaries across different *D. melanogaster* strains (see Methods), and genomic position and log2-normalized fold change in expression of heterochromatic genes in Su(var)2-10 GLKD versus control (shW) ovaries (based on RNA-seq). (B) Epigenetic profiles of heterochromatic Su(var)2-10 targets. From left to right: Heatmap of the normalized coverage and distribution of the H3K9me3 mark in control ovaries over the gene bodies and 2-kb flanking regions; heatmap of the percentage of RepeatMasker annotated regions within gene introns and 10-kb flanking regions; barplot of the fold change of steady-state RNA levels upon Su(var)2-10 GLKD (RNA-seq); barplot of the normalized H3K9me3 enrichment in control ovaries; and barplot of fold change in H3K9me3 levels in Su(var)2-10 GLKD (ChIP-seq). Data are averages from two biological replicates. (C) Epigenetic changes at heterochromatic genes down-regulated upon Su(var)2-10 GLKD. Shown are average profiles of normalized H3K9me3, H3K4me3, and Pol II ChIP signals across gene bodies and 2-kb flanking regions for Su(var)2-10 GLKD (orange) and control (green) ovaries. Shaded areas reflect standard errors. (D) Examples of transcriptomic and epigenetic changes at the heterochromatic gene *nAChRalpha4* upon Su(var)2-10 GLKD. UCSC browser tracks show the normalized read coverage of RNA-seq and H3K9me3, H3K4me3, and Pol II ChIP-seq data across the locus. Arrows mark direction of transcription. Top panel shows repetitive element annotations (RepeatMasker). Novel transcripts inferred from RNA-seq data are indicated by red bars. Red asterisk indicates ChIP peaks that appear in Su(var)2-10 GLKD libraries. Numbers on the right are the RPKM values of exonic regions (RNA-seq) or normalized ChIP/Input signal (ChIP-seq) in manually selected genomic intervals indicated by black bars. (E) Boxplot of log2-normalized RNA expression from sense exonic and antisense intronic regions of heterochromatic genes upon Su(var)2-10 GLKD. Data are averages of two biological replicates.

We found that heterochromatic genes are significantly over-represented among Su(var)2-10 targets (p<0.0001 Fisher’s exact test): 86 of the 228 heterochromatic genes expressed in the ovary displayed significant (≥2 fold) change in expression in cells depleted of Su(var)2-10 compared to control cells. However, in contrast to genes adjacent to TE insertions in euchromatin such as *jheh3* and *Frl/CG43986*, where loss of Su(var)2-10 led to transcriptional up-regulation, the majority of heterochromatic Su(var)2-10 targets (59 of the 86) were down-regulated upon Su(var)2-10 KD (Fig. 2A).

Heterochromatic genes typically have long introns enriched in TEs and other repetitive sequences. We found that heterochromatic genes regulated by Su(var)2-10 have high levels of H3K9me3 throughout their bodies (both over introns and exons) but not over their promoter regions, which are instead enriched in H3K4me2/3 and RNA polymerase II (Pol II), typical marks of active transcription (Fig. 2B, C). Thus, the allegedly repressive H3K9me3 mark is compatible with gene expression if present on the gene body and excluded from the promoter region. To understand the opposite effects of Su(var)2-10 on genes in eu- and heterochromatin, we analyzed the chromatin mark profile and expression of heterochromatic genes. Depletion of Su(var)2-10 changed the profiles of H3K9me3, H3K4me2/3, and Pol II over heterochromatic genes (Fig. 2C). Su(var)2-10 depletion led to a decrease in H3K9me3 signal on gene bodies and flanking regions. At the same time, marks of active transcription (H3K9me2/3 and Pol II) were also reduced at gene promoters, indicating that the observed change in steady-state RNA levels is due to transcriptional down-regulation. Taken together, our results indicate that Su(var)2-10-dependent deposition of H3K9me3 is required for the expression of genes residing in heterochromatin. This observation is in stark contrast with the canonical role of H3K9me3 as a repressive mark as in the case of TEs and euchromatic genes that have flanking transposon insertions (Fig. 1A).

To understand how loss of H3K9me3 interferes with the expression of heterochromatic genes, we carefully examined the RNA-seq profiles and enrichment of RNA polymerase II and H3K4me2/3 at the affected loci. We found that in several cases transcriptional down-regulation of the target gene upon Su(var)2-10 depletion was associated with the appearance of non-canonical transcripts from within introns or flanking regions, as illustrated for the *nAChRalpha4* gene (Fig. 2D). In wild-type ovaries, RNA-seq reads from *nAChRalpha4* mapped predominantly to exons, and few reads were generated from the long introns, which contain repetitive sequences. Upon Su(var)2-10 KD, the signals from exons were reduced, and signals from intronic transcripts, both sense and antisense with respect to the coding gene, were increased (Fig. 2D). The appearance of these non-canonical transcripts was associated with the presence of new H3K4me3 and Pol 11 peaks (Fig. 2D). The simultaneous appearance of intronic H3K4me2/3 signal marking active promoters and sense and antisense intronic transcripts indicates that loss of H3K9me3 upon Su(var)2-10 KD leads to activation of spurious transcription, potentially initiated by transposon promoters. Transcriptome analysis showed that intronic transcripts are non-canonical rather than inefficiently spliced pre-mRNA. Globally, there were increased levels of antisense intronic reads from the heterochromatic genes that were down-regulated upon germline depletion of Su(var)2-10 (Fig. 2E). Taken together, these results indicate that Su(var)2-10 is required for repression of spurious transcription likely initiated from TE promoters located in heterochromatin. T ranscription on either strand of long intronic regions or in flanking sequences of heterochromatic genes might interfere with promoter function.

Analysis of gene expression upon Su(var)2-10 GLKD revealed additional effects of H3K9me3 loss on heterochromatic genes. Loss of H3K9me3 correlated with activation of a cryptic promoter in an intron of the *Nipped-B* gene that led to the appearance of a new mRNA isoform (Fig. 3A). This new isoform is missing 14 coding exons of the canonical *Nipped-B* transcript and has one new exon, which shares homology with the *gypsy* retrotransposon. Expression of the full-length *Nipped-B* isoform is only moderately affected by Su(var)2-10 depletion, but expression of the truncated protein from the new mRNA isoform might cause dominant negative effects and interfere with proper protein function. In the case of *unc-13*, Su(var)2-10 GLKD and associated H3K9me3 loss resulted in the up-regulation of a longer isoform from an upstream transcription start site (TSS) adjacent to an *Invader* transposon and in reduced activity of the canonical TSS (Fig. 3B). Ectopic transcriptional activation can affect genes at long distances as evident for the gene *CG10417*, which encodes a conserved protein phosphatase (Fig. 3C). H3K9me3 loss and transcriptional up-regulation near a TE-rich region about 5 kb upstream of the canonical *CG10417* promoter resulted in the emergence of a long fusion transcript between the upstream region and the protein-coding gene. As was observed for *unc-13*, up-regulation of transcription from the upstream TSS was associated with reduced H3K4me3 at the canonical promoter and with repression of the normal mRNA isoform (Fig. 3C). Collectively, these examples show that loss of H3K9me3 causes activation of TE transcription within introns or in regions adjacent to host genes and interferes with the activity of gene promoters, leading to reduced gene expression and/or appearance of new mRNA isoforms encoding novel proteins. Overall, our results indicate that genes positioned in heterochromatin have very different properties than their euchromatin counterparts: Genes in heterochromatin require high levels of H3K9me3 for proper expression and canonical isoform selection.

**Figure 3.**
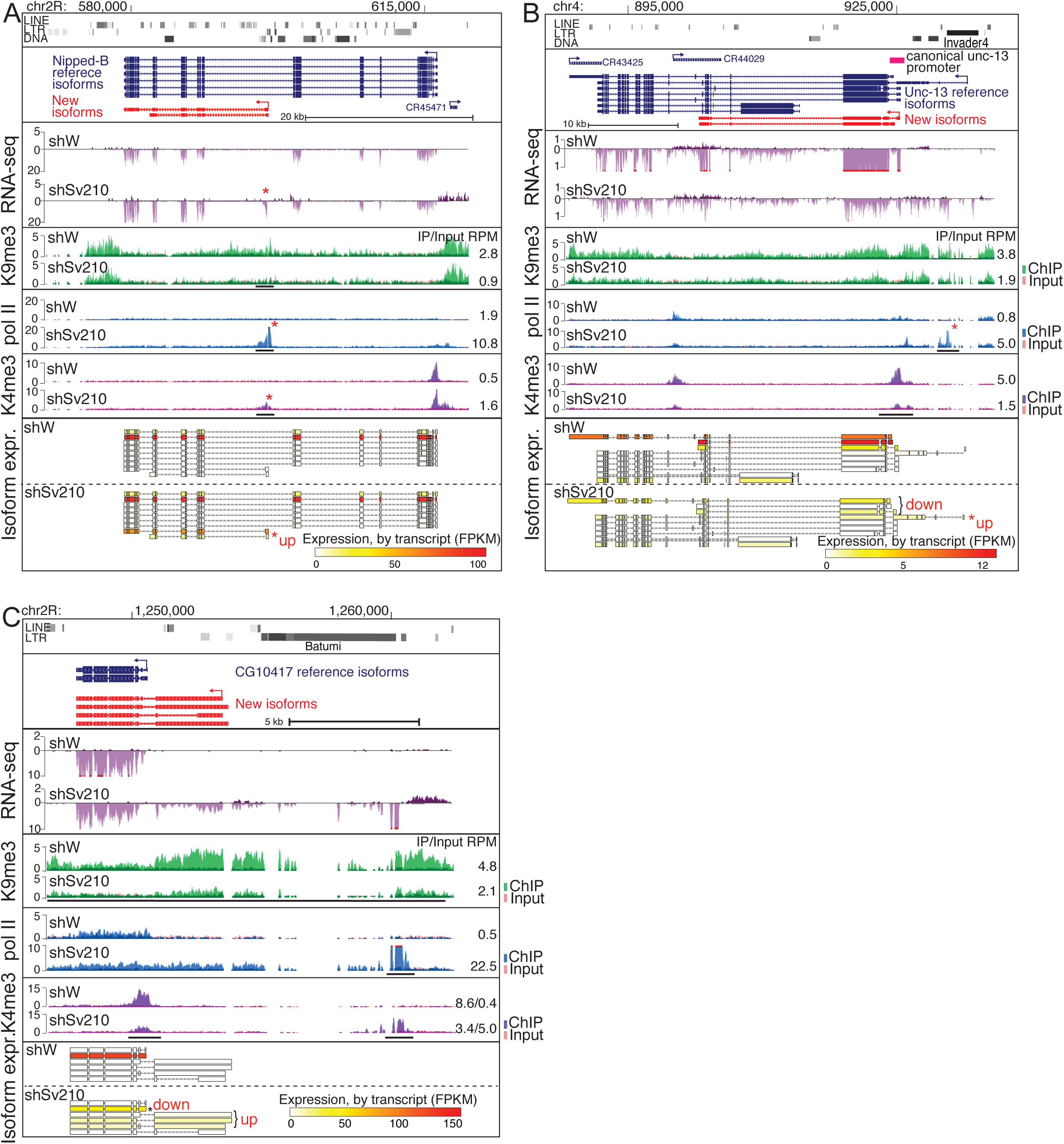
H3K9me3 loss induces transcriptional interference and aberrant isoform expression. (A) Su(var)2-10 loss leads to emergence of a novel, truncated *Nipped-B* isoform. UCSC browser tracks show the normalized read coverage of RNA-seq and H3K9me3, H3K4me3, and Pol II ChIP-seq experiments across the locus. New isoforms (see Methods) are shown in red. Arrows mark direction of transcription. Red asterisks mark differentially up-regulated novel region and corresponding peaks of H3K4me3 and Pol II. Bottom panels show the relative expression of detected isoforms. (B, C) Su(var)2-10 loss causes activation of upstream alternative TSSs and repression of canonical TSSs in *unc-13* and *CG10417* loci. UCSC browser tracks show the RNA-seq and ChIP-seq signal in control and Su(var)2-10 GLKD ovaries. In all panels, numbers on the right to ChIP-seq tracks are the normalized ChIP/Input signal (ChIP-seq) in manually selected genomic intervals indicated by black bars.

### Su(var)2-10-dependent installation of H3K9me3 restricts expression of tissue-specific genes

The presence of TE sequences that are targeted by Su(var)2-10 in close proximity to either hetero- or euchromatic genes explains the effect of Su(var)2-10 on such genes. However, a number of euchromatic genes repressed by Su(var)2-10 do not have proximal TE insertions in the reference genome, and we did not find evidence of non-reference insertions using TIDAL analysis. Some of these genes might be secondary targets that have altered expression upon depletion of Su(var)2-10 as a result of changes in expression of primary target genes. However, a subset of euchromatic Su(var)2-10 targets have local H3K9me3 peaks and examination of additional H3K9me3 ChIP-seq datasets from previous studies (LeThomas et al., 2013; Muerdter et al., 2013; Yu et al., 2015, see Methods) showed that a number of these H3K9me3 peaks are present in different *D. melanogaster* strains (Fig. 4A). Thus, these genes may represent Su(var)2-10 targets that are repressed though H3K9me3 deposition independently of TEs.

**Figure 4.**
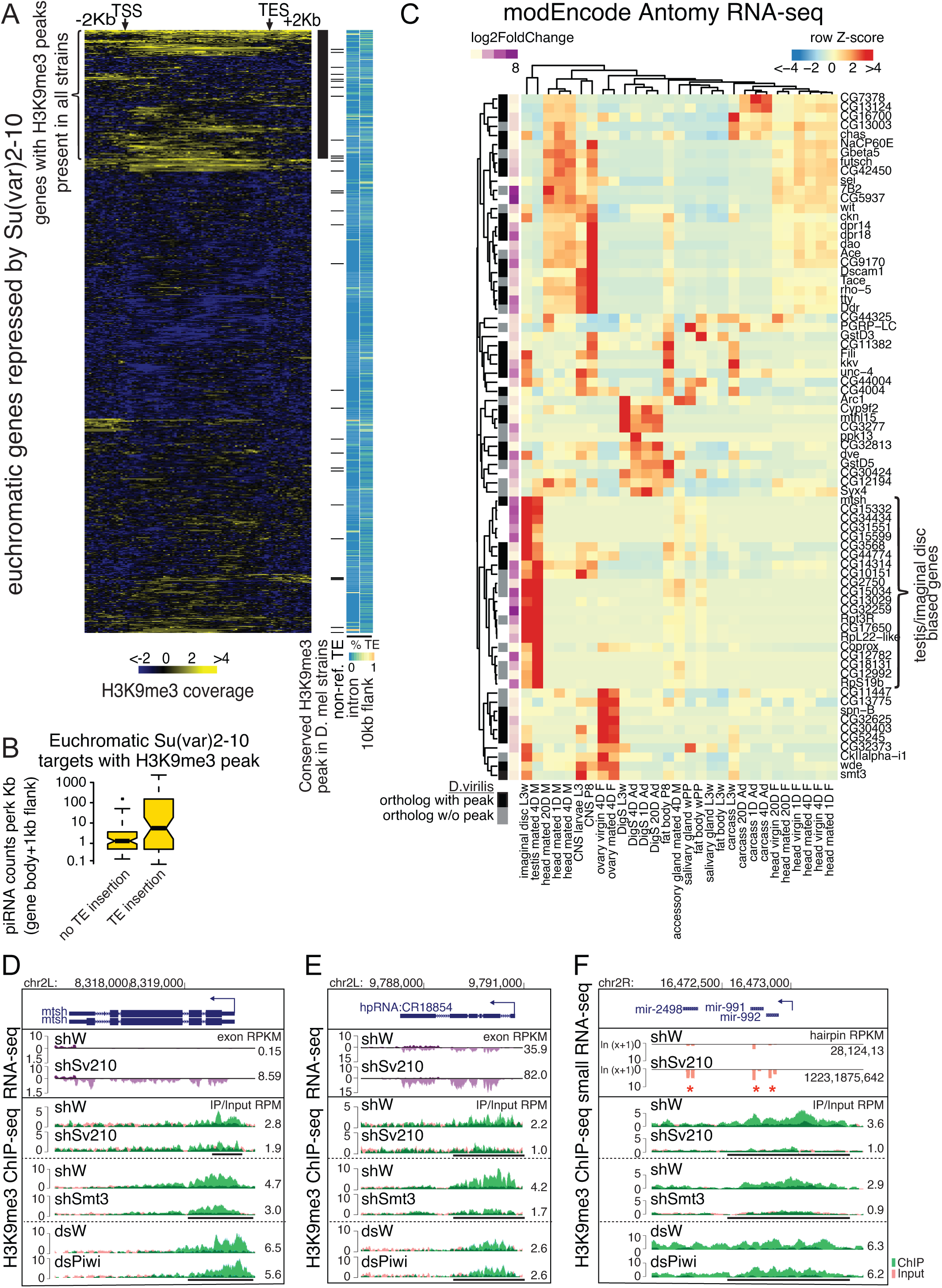
Loss of H3K9me3 islands over euchromatic genes leads to ectopic gene expression. (A) A subset of euchromatic genes repressed by Su(var)2-10 have discrete H3K9me3 peaks consistent between different strains of *Drosophila.* Heatmap of the normalized coverage of H3K9me3 mark in control ovaries over the gene bodies and 2-kb flanking regions of Su(var)2-10-repressed euchromatic genes. Side bars indicate, for each gene, whether H3K9me3 peaks are present in ovaries of different *D. melanogaster* strains (see Methods), whether non-reference TE insertions are present within gene bodies and 1-kb flanking regions, the percentage of RepeatMasker annotated regions within gene introns, and the percentage of RepeatMasker annotated regions within 10-kb flanking regions. (B) Boxplot of the number of 23-29-nt RNAs that can be aligned to Su(var)2-10/H3K9me3 repressed genes and 1-kb flanking regions localized near TEs or in TE-free regions. (C) Heatmap of z-scaled expression levels of euchromatic genes localized in TE-free regions that are de-repressed upon Su(var)2-10 GLKD and that harbor H3K9me3 peaks. Tissue expression data was retrieved from the modEncode Anatomy database. The sidebar indicates whether genes have orthologues with conserved H3K9me3 peaks in *D. virilis.* (D-F) UCSC browser tracks show the normalized read coverage of RNA-seq and H3K9me3 ChIP-seq signal for the *mtsh* locus, E) the endo-siRNA locus *CG18854*, and F) the small RNA-seq and H3K9me3 ChIP-seq signal for the *mir-991/992/2498* locus in indicated knockdown and corresponding control ovaries. Numbers on the right in panels D and E are the RPKM-normalized mRNA expression levels. Numbers on the right in panel F are RPM-normalized microRNA read counts for each hairpin (small RNA-seq). Normalized ChIP/Input signals (ChIP-seq) in manually selected genomic intervals are indicated by black bars.

It is well established that H3K9me3 deposition at transposons is mediated by the piRNA pathway. To further confirm that the putative TE-independent H3K9me3 peaks on euchromatic Su(var)2-10 targets are deposited independently of the piRNA pathway, we analyzed whether piRNAs map within the gene bodies and flanking regions of these genes using small RNA libraries from ovaries of the same strain. In contrast to genes with TE insertions inside or proximal to the gene, regions flanking Su(var)2-10 targets with no TE insertions were devoid of piRNA-sized reads (Fig. 4B).

We then asked if H3K9me3 peaks at Su(var)2-10 repressed genes in euchromatin are conserved in evolution. Analysis of available H3K9me3 ChIP-seq data (LeThomas et al., 2014) indicated that more than half (32 out of 58) of the *D. melanogaster* genes suppressed by Su(var)2-10 that do not have adjacent TE insertions have *D. virilis* homologs marked by H3K9me3 peaks. *D. virilis* diverged from *D. melanogaster* more than 45 million years ago (Hedges and Kumar, 2009), and it is unlikely that active TEs are conserved between the two species. Therefore, conserved H3K9me3 signals on homologous genes of the two species are indicative of TE-independent deposition. Taken together these data suggest that Su(var)2-10 is recruited to a number of genes in euchromatin in a transposon-independent fashion to restrict their expression.

Genes that are regulated by Su(var)2-10 due to proximal TE insertions are not enriched in certain pathways (data not shown). However, we found that many targets of TE-independent repression by Su(var)2-10/H3K9me3 have expression patterns that are biased to specific tissues such as the central nervous system or head, larval imaginal discs, or the testis (Fig. 4C). These genes were typically not expressed in control ovaries, but depletion of Su(var)2-10 led to ectopic expression in the female germline. For example, the testis-specific gene *mtsh* (which encodes a factor involved in male meiosis) is repressed in the ovary and marked by a strong H3K9me3 peak at its putative promoter region that depends on Su(var)2-10 and SUMO, but not on Piwi (Fig. 4D). Su(var)2-10 GLKD led to loss of the H3K9me3 peak and ectopic expression of *mtsh.* H3K9me3 regulation is not limited to protein coding-genes: Transcription from two well-characterized testis-biased endogenous hp-siRNA loci (Wen et al., 2015) is repressed through Su(var)2-10-dependent H3K9me3 deposition (Fig. 4E; Fig. S1). The putative promoter region of the testis-specific microRNA (miRNA) cluster *mir-992/991/2498* (Mohammed et al., 2014) is also marked by a strong H3K9me3 peak, and expression of these miRNAs is repressed in the ovary. Su(var)2-10 GLKD led to loss of the H3K9me3 peak and concomitant ectopic expression of the three microRNAs as shown by analysis of small RNA-seq libraries (Fig. 4F). Thus, Su(var)2-10 plays a role in maintaining appropriate female-specific expression patterns in germline cells by repression of testis-expressed genes. More generally, Su(var)2-10 prevents inappropriate expression of genes that are normally expressed tissue specifically through a mechanism that is independent of transposon suppression.

### Su(var)2-10 represses a set of genes involved in heterochromatin formation

Su(var)2-10 also represses several genes that encode proteins involved in heterochromatin formation and maintenance (Fig. 5A). Heterochromatin is established and maintained though an interplay between histone mark writer complexes that deposit repressive histone marks such as H3K9me3 and reader complexes that recognize repressive marks and recruit downstream heterochromatin components. Su(var)2-10 GLKD led to increased expression of wde. This gene encodes a conserved co-factor of the essential H3K9 methyltransferase SetDB1/Eggless, which is responsible for the installation of the H3K9me3 mark at a large proportion of heterochromatic domains (Clough et al., 2007; Koch et al., 2009; Rangan et al., 2011; Seum et al., 2007; Timms et al., 2016; Tzeng et al., 2007; Wang et al., 2003). Su(var)2-10 also represses expression of two recently identified proteins that interact with the protein HP1, which is critical for heterochromatin packaging: Sov and CG30403. Sov is a suppressor of positional effect variegation and plays a role in recruitment of HP1 to chromatin (Alekseyenko et al., 2014; Jankovics et al., 2018). To further investigate the role of Sov in heterochromatin formation, we tested the effect of *sov* GLKD on the genome-wide distribution of the H3K9me3 mark in the ovary using ChIP-seq. H3K9me3 levels decreased globally upon depletion of *sov* (Fig. S2A, B), indicating that Sov is required for H3K9me3 deposition, explaining its role in HP1a recruitment.

**Figure 5.**
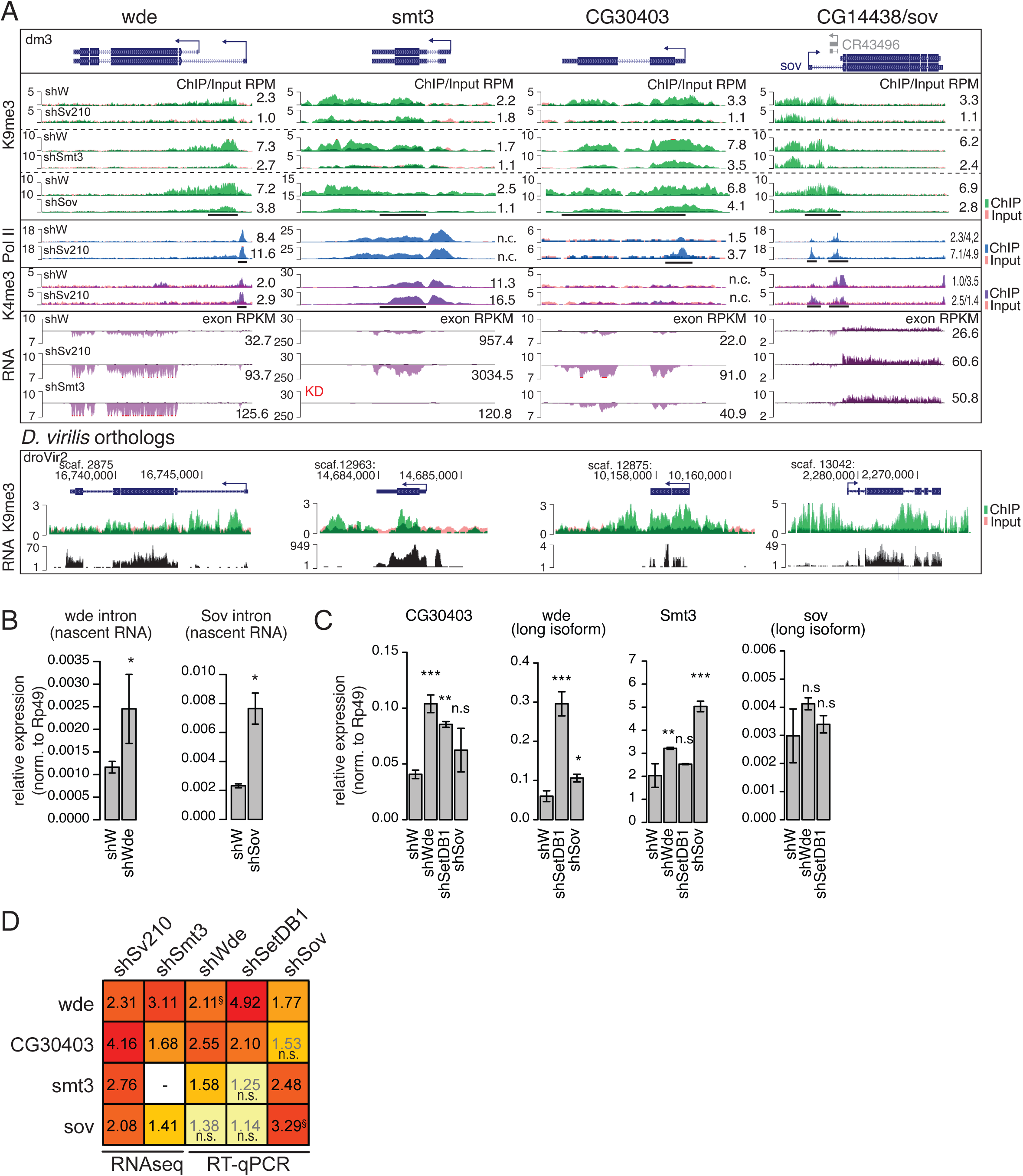
The H3K9me3 mark regulates factors involved in heterochromatin formation. (A) UCSC browser tracks of the normalized signal from RNA-seq and ChIP-seq in GLKD and control ovaries for four factors involved in heterochromatin regulation. Bottom panel shows conservation of the H3K9me3 peaks at orthologous loci in *D. virilis.* Numbers on the right are RPKM-normalized mRNA expression levels (eXpress) or the normalized ChIP/Input signal (ChIP-seq) in manually selected genomic intervals indicated by black bars. Data are representative of two biological replicates. (B) Barplot of the relative expression of *wde* and *sov* nascent transcripts (RT-qPCR) in ovaries of controls (shW) and flies depleted of Wde or Sov proteins. Fold changes in the nascent transcripts were measured using intronic primers. Error bars show standard deviations from three biological replicates. The asterisks indicate significant differences compared to shW, p<0.05 (Student’s two tailed t-test). (C) Barplot of the relative gene expression of indicated genes (RT-qPCR) in control (shW) ovaries and upon indicated GLKD by shRNA. Error bars show standard deviations from three biological replicates. Asterisks indicate significant difference compared to the shW samples, *p<0.05, **p<0.01, ***p<0.001; “n.s.” indicates not significant (Student’s two tailed t-test). (D) Heatmaps of all-versus-all summaries of heterochromatin factor fold changes upon GLKD compared to corresponding shW controls. Values are fold changes in knockdown vs. corresponding shW controls determined by DESeq2 for RNA-seq derived data (from experiments shown in panel A) or average relative expression levels (experiments shown in panels B and C). “n.s.” denotes values that were not statistically significant according to DESeq2 analysis (RNA-seq data) or 2-tailed Student’s t-test (RT-qPCR data). §For regulation of *sov* and *wde* upon their corresponding mRNA knockdown, shown are fold changes in the nascent transcripts measured by intronic primers (see main text and panel B).

*CG30403* encodes a DNA-binding protein that emerged as an HP1a interactor in a proteomic screen (Alekseyenko et al., 2014). We confirmed that CG30403 interactrs with HP1 by co-immunoprecipitation from *Drosophila* S2 cell lysate (Fig. S2C). We were unable to achieve efficient depletion of CG30403 using an shRNA prohibiting further investigation of its function. Finally, Su(var)2-10 GLKD led to up-regulation of the *smt3* gene encoding the *Drosophila* SUMO (Small Ubiquitin-like Modifier) homolog. SUMO has been implicated in various aspects of heterochromatin formation and transcriptional repression in organisms ranging from yeast to human, and we showed that SUMO is required for H3K9me3 deposition downstream of Su(var)2-10 and for TE repression in the *Drosophila* ovary in the accompanying manuscript (Ninova et al., accompanying manuscript).

The four genes encoding proteins involved in heterochromatin formation, wde, sov, *CG30403*, and *smt3*, all have H3K9me3 peaks that are deposited in a manner dependent on both Su(var)2-10 and SUMO (Fig. 5A). In all four cases, the peaks were conserved at the orthologous genes in *D. virilis* (Fig. 5A). Furthermore, we found no evidence of transposons adjacent to *wde, smt3*, or *CG30403* (from DNA-seq data and genomic PCR, data not shown) or for piRNAs aligning to these regions, strongly suggesting that these genes are direct, TE-independent targets of H3K9me3 deposition through Su(var)2-10/SUMO. Examination of the chromatin environment around these loci suggested that in all cases, increased expression upon Su(var)2-10 GLKD is a consequence of H3K9me3 loss and transcriptional up-regulation (Fig. 5A). For instance, *wde* is expressed from two alternative promoters located on the opposite sides of a prominent H3K9me3 island. Loss of H3K9me3 upon Su(var)2-10 depletion correlated with an increased use of the upstream promoter and an overall increase in *wde* expression as evidenced by RNA-seq and Pol II and H3K4me3 ChIP-seq (Fig. 5A). Similarly, Su(var)2-10 and SUMO were required for the presence of H3K9me3 islands and for the repression of sov, *CG30403*, and *smt3* (Fig. 5A). In the case of sov, we observed increased transcriptional activity from an upstream TSS upon Su(var)2-10 depletion, whereas *CG30403* and *smt3* up-regulation occurred due to enhanced transcription from single promoters. Taken together, our data suggests that Su(var)2-10 is involved in H3K9me3-mediated repression of four genes that encode proteins involved in heterochromatin formation.

### Expression of genes encoding heterochromatin proteins is regulated by a negative feedback loop

The observation that genes encoding heterochromatin proteins are themselves repressed by local H3K9me3-rich chromatin islands suggests that their expression might be controlled by a negative feedback loop in which gene activity is repressed by its own product. To test if such negative feedback occurs, we asked if depletion of the gene product affected the corresponding gene expression. To this end, we took advantage of the facts that RNAi destroys mRNAs in the cytoplasm causing protein depletion and that the abundance of nascent (non-spliced) pre-mRNA can by analyzed by RT-PCR to give a proxy of the level of transcription. We found that GLKD of *wde* and *sov* led to increases in abundance of their respective pre-mRNAs, confirming that expression of these genes is auto-regulated (Fig. 5B). *sov* auto-regulation was further evident on the level of chromatin, as *sov* GLKD resulted in a decrease in the H3K9me3 ChIP-seq signal at the *sov* promoter (Fig. 5A). We were unable to test *CG30403* and *smt3* auto-regulation due to poor knockdown efficiency and lack of utilizable introns, respectively.

In the accompanying manuscript, we show that tethering of Su(var)2-10 to chromatin induces SUMO-dependent recruitment of the SetDB1/Wde histone methyltransferase complex, resulting in H3K9me3 deposition and transcriptional silencing (Ninova et al., accompanying manuscript). Wde, *SUMO(smt3)*, CG30403, and Sov might function independently to mediate heterochromatin formation or may participate in one common pathway that regulates expression of all four genes. To discriminate between these two possibilities, we explored the dependence of these genes on each other as well as on the effector methyltransferase SetDB1. Analysis of RNA-seq data from ovaries in which *smt3* expression was inhibited in germ cells showed that wde, *CG30403*, and *sov* are up-regulated upon *smt3* loss (Fig. 5A). Expression of *CG30403* and *wde* was increased >2 fold upon GLKD of *SetDBI* and wde, and slightly in *sov* GLKD ovaries (Fig. 5C, D). In contrast, *smt3* and *sov* transcript levels showed modest or no significant increase upon *SetDBI* and *wde* depletion. Taken together, these results indicate that Su(var)2-10, SetDB1/Wde, and Smt3 act in the same pathway to confer H3K9me3 deposition and transcriptional repression at the *wde* and *CG30403* loci. The observation that repression of *smt3* and *sov* is not strongly impacted by depletion of SetDB1 and Wde implies existence of a distinct SUMO/Su(var)2-10/Sov-dependent silencing pathway that does not require the SetDB1/Wde complex.

## Discussion

### Su(var)2-10 and H3K9me3 play important roles in the regulation of gene expression

Histone modifications play an essential role in the control of gene expression and in the nuclear packaging of the genome. H3K9me3 is a conserved hallmark of constitutive heterochromatin from yeast to human and is generally associated with gene repression. We identified Su(var)2-10 as a crucial factor required for deposition of the H3K9me3 mark in large heterochromatic domains of the *D. melanogaster* genome, such as pericentromeric regions and the 4^th^ chromosome, as well as at discrete islands in euchromatin. In the accompanying manuscript, we demonstrate that Su(var)2-10 localization to chromatin induces strong transcriptional silencing and that Su(var)2-10 acts in a SUMO-dependent manner to recruit the histone-methyltransferase complex composed of SetDB1 and Wde to chromatin (Ninova et al., accompanying manuscript)

Su(var)2-10-dependent H3K9me3 deposition is crucial to ensure proper TE silencing in the germ cells of *Drosophila* (Ninova et al., accompanying manuscript). Here we showed that in addition to effects on transposon silencing Su(var)2-10 and H3K9me3 influence regulation of gene expression of protein-coding genes. Su(var)2-10-dependent H3K9me3 deposition on TEs impacts expression of genes located in heterochromatin and expression of euchromatic genes adjacent to TE insertions. Su(var)2-10 is also involved in H3K9me3 deposition on host genes independently of TEs. This process is essential for suppression of ectopic expression of tissue-specific genes, thereby conferring correct cell type identity.

### Epigenetic effects of transposons influence host gene expression

Approximately half of the human genome is comprised of transposon sequences, and the TE fraction is as high as 90% in several plant species (Lander et al., 2001; Sabot et al., 2005; Schnable et al., 2009). It is estimated that in humans one new transposon insertion per generation is propagated to offspring (Kazazian, 2004). The numbers of somatic TE insertions, although difficult to detect, are likely much higher (Kazazian and Jr, 2011). Thus, transposon activity is a major source of genetic variation, and this variation can occur on a very short time scale. The effects of transposons on the host transcriptome have been the subject of many studies beginning with the pioneering work by Barbara McClintock, who identified “control” elements that regulate gene expression before genome compositions were known (McClintock, 1950). Subsequent work in many organisms has shown that transposons can disrupt gene expression by inserting into coding gene regions or into or close to cis-regulatory sequences such as promoters and enhancers (Chuong et al., 2017). In fact, many transposons tend to insert close to promoters and enhancers (Chuong et al., 2017; Galvan et al., 2009; de Jong et al., 2014; LaFave et al., 2014; Liao et al., 2000; Wu et al., 2003). Notably, TE insertions are not always disruptive: Insertions into non-coding regions can bring new regulatory elements that change gene expression patterns resulting in increased fitness (Feschotte, 2008). Instances of positive selection for TE insertions are well documented in *Drosophila* (González et al., 2008; Maside et al., 2002; Schlenke and Begun, 2004). Transposon-derived promoters also drive expression of numerous mouse and human genes, suggesting that TE insertions can be co-opted into gene regulatory pathways (Chuong et al., 2017; Jordan et al., 2003; Nigumann et al., 2002; Peaston et al., 2004).

In addition to direct changes in the DNA sequence, TE insertions may also introduce local epigenetic effects (Slotkin and Martienssen, 2007). Active transposons are transcriptionally silenced by H3K9 trimethylation and/or DNA methylation. The H3K9me3 mark can spread several kilobases outside the transposon region (Lee and Karpen, 2017; Pezic et al., 2014; Sentmanat and Elgin, 2012; Sienski et al., 2012). Therefore, H3K9me3 deposition due to TE integration may affect adjacent cis-regulatory elements of host genes, interfering with their normal expression. Indeed, TE insertions with high levels of H3K9me3 are strongly selected against, supporting a model that TEs can alter expression of host genes through epigenetic changes (Lee and Karpen, 2017).

The finding that Su(var)2-10 is responsible for deposition of H3K9me3 on TE bodies and flanking sequences allows us to separate the effect of direct damage to *cis*-regulatory elements from the effect on chromatin. We found evidence that TE insertions can lead to H3K9me3-dependent changes in gene expression. As shown for the *jheh3* and *frl* loci (Fig. 1B), loss of the TE-associated H3K9me3 peak upon Su(var)2-10 GLKD resulted in up-regulation of expression of both genes. Notably, it was previously shown that the *BARI* insertion at the *jheh* locus was under positive selection in *D. melanogaster* populations (Gonzalez et al., 2009), indicating that transposon-induced epigenetic repression of *jheh3* is beneficial for the host. Overall, our results suggest that transposons can rewire gene regulatory networks on a short time scale due not only to alterations in the genome but also to effects on chromatin. Euchromatic H3K9me3 peaks due to TE insertions are widespread in *Drosophila* (Lee, 2015; Lee and Karpen, 2017; Sienski et al., 2012), indicating that gene expression modulation by proximal TE insertion may be a common mode of introducing gene regulatory variation. Our results suggest that differential gene expression between *Drosophila* strains might be partly due to polymorphic TE insertions. New TE insertions during development reportedly generate genomic diversity between different cell types in human and mouse with implications for tumorigenesis and brain development (Faulkner and Garcia-Perez, 2017). Future studies are required to elicit the epigenetic effects of somatic TE insertions on gene regulatory networks.

### Genes located in heterochromatin require H3K9me3 for proper expression

In *Drosophila*, heterochromatin domains include nearly 30% of the genome (Hoskins et al., 2002, 2007). Even though these regions are relatively gene poor, heterochromatin hosts several hundred protein-coding genes. Studies of chromosomal rearrangements suggested that heterochromatic localization is in fact required for proper expression of genes in heterochromatin (Yasuhara and Wakimoto, 2006). For example, translocation of the *light* gene, which is normally localized in the pericentromeric region of chromosome 2L, to euchromatin results in reduced and variegated expression (Wakimoto et al., 1990). However, the molecular mechanism of the positive effect of the heterochromatin environment on expression is not completely understood.

Consistent with previous studies, our results indicate that despite residing in H3K9me3-enriched loci, certain heterochromatic genes are not silenced (Riddle et al., 2011, 2012). Unlike genes adjacent to TE insertions in euchromatin, which are suppressed by Su(var)2-10-dependent deposition of H3K9me3 mark, a substantial fraction of expressed heterochromatic genes require Su(var)2-10 and the H3K9me3 mark for their expression (Fig. 2, 3). Thus, our results show that the allegedly repressive H3K9me3 mark does not interfere with transcription but is actually necessary for the expression of genes in heterochromatin.

How can the same chromatin mark lead to repression of genes in euchromatin and activation in heterochromatin? The H3K9me3 mark is present over the gene bodies and regions flanking heterochromatic genes; however, promoters of these loci are depleted of H3K9me3 and instead carry typical marks of active promoters such as H3K4me3 and Pol II occupancy. Thus, H3K9me3 over gene bodies appears to be compatible with transcription. Depletion of H3K9me3 upon Su(var)2-10 GLKD correlated with increased levels of intronic RNAs and the appearance of H3K4me2/3 and Pol II signals in introns, indicating up-regulation of non-canonical transcripts originating from within host-gene introns. One possible source of such transcripts is the activation of transposon promoters that are highly abundant within introns and flanking sequences of heterochromatic genes. For instance, activation of transposon promoters in the *unc-13* and *CG10417* loci reduced the expression from the canonical gene promoters (Fig. 3B, C). We propose that transcription from transposon promoters located in introns and flanking sequences interferes with proper gene expression through transcriptional interference (Shearwin et al., 2005).

Loss of the H3K9me3 mark also disrupted normal isoform regulation of heterochromatic genes, as we observed both truncated and extended mRNA isoforms with coding potential distinct from the canonical gene mRNA upon depletion of Su(var)2-10 (Fig. 3). Activation of cryptic promoters upon H3K9me3 loss might disrupt proper gene expression through mechanisms other than simple reduction in canonical mRNA output. For example, expression of novel proteins from non-canonical mRNA isoforms might cause dominant negative effects. It should be noted that not all Su(var)2-10-dependent heterochromatic genes that lose H3K9me3 upon Su(var)2-10 GLKD show signs of non-canonical transcription activation. This indicates that H3K9me3 might have other functions in heterochromatic gene activation. For example, compaction of heterochromatin by H3K9me3-associated HP1 might bring distant enhancers of heterochromatic genes into physical proximity of promoters to activate expression. Overall, our results combined with previous studies indicate that, in contrast to euchromatic genes, genes positioned in heterochromatin require high H3K9me3 levels for proper expression and isoform selection.

### H3K9me3 in euchromatin restricts gene expression to correct cell lineages

Although the H3K9me3 mark is usually associated with constitutive heterochromatin and TE repression, discrete Su(var)2-10-dependent H3K9me3 peaks are present at a number of euchromatic genes. We found no evidence of TEs in the vicinity of some of these euchromatic genes, and their H3K9me3-based repression was conserved between D. *melanogaster* and D. *virilis* – two species that separated more than 45 million years ago and have no common transposon insertions (Fig. 4A, B). We found that expression of many of the genes marked by TE-independent H3K9me3 is restricted to specific tissues and cell types such testis, digestive system, or central nervous system and that the loss of H3K9me3 upon Su(var)2-10 depletion resulted in ectopic expression in the female germline. Our finding that H3K9me3 represses testis-specific genes is in line with a recent report that SetDB1 depletion in the female germline was associated with loss of the H3K9me3 mark and results in mis-expression of male-specific genes (Smolko et al., 2018). Thus, gene repression through H3K9me3 plays a crucial role in maintaining proper expression patterns in female germline. H3K9me3, the SetDB1 methyltransferase, and the SUMO pathway are also implicated in lineage-specific gene expression and cell fate commitment in mammalian systems (Becker et al., 2016; Cheloufi et al., 2015; Cossec et al., 2018; Ivanov et al., 2007; Wang et al., 2018; Yang et al., 2015). Collectively, these data suggest that a TE-independent H3K9me3 deposition via the SUMO-SetDB1 pathway plays an important and evolutionarily conserved role in restricting gene expression to proper cell types and lineages.

### Negative feedback regulation of ‘heterochromatin sensors’ ensures chromatin homeostasis in the cell

We found that in addition to tissue-restricted genes, SUMO- and Su(var)2-10-dependent H3K9me3 repression is also necessary for the regulation of several factors involved in heterochromatin formation and maintenance, such as SUMO *(smt3)*, the SetDB1-cofactor Wde, and two partners of HP1, Sov and CG30403. Wde is the homologue of the mammalian MCAF1/ATF7IP; this factor is required for nuclear localization and stability of SetDB1 (Koch et al., 2009; Timms et al., 2016) and promotes its methyltransferase activity (Wang et al., 2003). *Drosophila* Wde also associates with SetDB1, and germline depletion of Wde results in a phenotype similar to depletion of SetDB1, supporting the role of Wde as a conserved SetDB1 co-factor (Koch et al., 2009; Smolko et al., 2018). Our data in *Drosophila* and studies in mammals suggest that SUMO is involved in the recruitment of SetDB1/Wde complex to its chromatin targets (Ninova et al., accompanying manuscript, Ivanov et al., 2007). HP1 is a critical reader of the H3K9me3 mark, and HP1 multimerization is responsible for the structural properties of heterochromatin (Canzio et al., 2011; Hiragami-Hamada et al., 2016; Larson et al., 2017; Strom et al., 2017). Furthermore, HP1 is a hub that interacts with many other heterochromatin proteins (Eissenberg and Elgin, 2014). Both Sov and CG30403 interact with HP1, and Sov is critical for heterochromatin maintenance (Jankovics et al., 2018, Fig. S1).

The genes encoding Wde, SUMO, Sov, and CG30403 reside in euchromatin and are repressed by local H3K9me3 marks. In contrast to tissue-restricted genes, which are often completely repressed by Su(var)2-10 in female germline, expression of these factors is not silenced. However loss of H3K9me3 upon Su(var)2-10 depletion leads to their transcriptional up-regulation (Fig. 5A). Our results indicate that these four genes encoding proteins involved in heterochromatin formation are part of a negative feedback mechanism that controls heterochromatin formation. Negative feedback in biological circuits often maintains protein levels within a certain range, providing homeostatic regulation. We propose that SUMO-dependent repression of heterochromatin proteins provides such homeostatic regulation to maintain the proper ratio and boundaries of hetero- and euchromatin. According to our model (Fig. 6), specific genes, such as wde, act as sensors of overall H3K9me3 levels. Insufficient levels of H3K9 methylation lead to elevated expression of the sensor genes due to decreased levels of H3K9me3 at their promoters. The consequential increased sensor expression will enhance H3K9me3 deposition and heterochromatin formation throughout the genome. This in turn will result in repression of sensor genes, ensuring that H3K9me3 is restricted to proper genomic domains and does not spread to euchromatic regions that should remain active.

**Figure 6.**
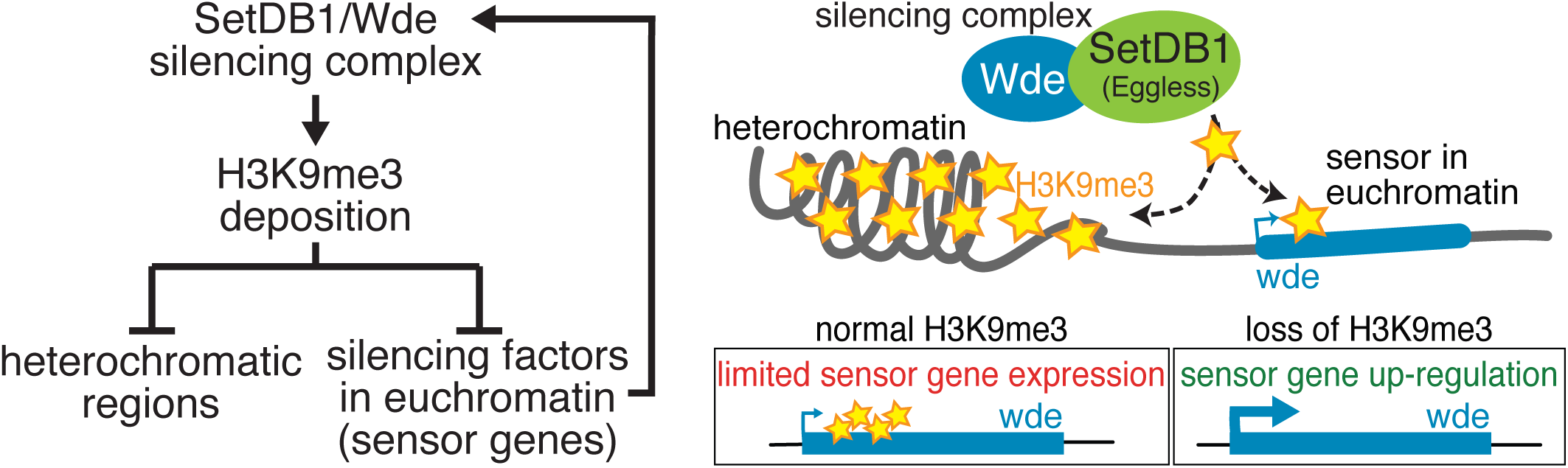
Model of how euchromatic sensor genes regulate cellular H3K9me3 levels. Sensor genes such as *wde* encode essential components of the heterochromatin maintenance machinery. These genes reside in canonical euchromatin but are themselves repressed by H3K9me3 located over their gene bodies. Loss of H3K9me3 over these sensors leads to up-regulation, and production of these proteins, such as Wde, facilitating re-establishment of normal levels of H3K9me3 throughout the genone and at sensor genes themselves, thereby limiting expression and further silencing activity of the sensors.

Controlling heterochromatin spreading is critical for gene expression as illustrated by the phenomenon of position effect variegation. Chromosomal rearrangements that bring euchromatic genes in proximity to heterochromatin cause their repression by spreading of heterochromatin (Elgin and Reuter, 2013). Our findings (Fig. 2, 3) and previous reports (reviewed in Yasuhara and Wakimoto, 2006) demonstrated that loss of the heterochromatic environment interferes with expression of genes located in regions that would normally be heterochromatic. Thus, maintaining the correct ratio and boundaries between chromatin domains via a negative feedback mechanism is crucial for proper gene expression in both domains. A reminiscent negative feedback loop was identified in yeast: Genes essential for heterochromatin assembly such as the H3K9 methyltransferase *clr4* are suppressed by H3K9me3 to restrict ectopic spreading of silencing chromatin (Wang et al., 2015). In mammals, genes encoding proteins from the KRAB-ZFP family of transcriptional repressors reside in H3K9me3- and HP1-enriched loci (Frietze et al., 2010; O’Geen et al., 2007; Vogel et al., 2006). Collectively, these studies suggest that auto-regulation of heterochromatin effectors is a conserved mode of chromatin regulation, although the specific genes involved in the feedback mechanism differ between different organisms. In the future, it will be important to dissect the precise network architecture of heterochromatin regulation. As heterochromatin formation and maintenance was reported to be disrupted in cancer and during aging this mechanism might be a promising target of therapeutic interventions.

### Molecular mechanism of Su(var)2-10 recruitment to genomic targets

Studies in various systems have identified diverse mechanisms for recruitment of H3K9me3 writer enzymes to target loci. In the case of TE repression in germ cells, piRNAs bound to nuclear Piwi proteins serve as sequence-specific guides that bind complementary nascent transcripts (LeThomas et al., 2013; Rozhkov et al., 2013; Sienski et al., 2012) that in turn recruit Su(var)2-10, which induces H3K9me3 deposition in a SetDB1-dependent manner (Ninova et al., accompanying manuscript). The majority of Piwi-bound piRNAs are complementary to transposon mRNAs (Brennecke et al., 2007), ensuring that Su(var)2-10 and the SetDB1/Wde complex is recruited to transcribed TE copies regardless of their genomic position. H3K9me3 targeted to TE insertions can spread to the adjacent loci encoding host genes (Lee, 2015; Lee and Karpen, 2017; Sentmanat and Elgin, 2012). Our study demonstrates another mechanism of writer recruitment: We found that Su(var)2-10 identifies non-transposon targets in a piRNA-independent fashion (Fig. 4A) in agreement with a broader function of Su(var)2-10 in development (Hari et al., 2001). The observation that H3K9me3 peaks at homologous euchromatic genes are also present in the distantly related *D. virilis* points to a conserved mechanism for H3K9me3 deposition involved in host-gene regulation.

The molecular mechanism of piRNA-independent recruitment of Su(var)2-10 remains to be explored. Su(var)2-10 has a putative DNA binding SAP domain that might be sufficient for its binding to DNA (Aravind and Koonin, 2000). However, motif enrichment analysis failed to identify a common sequence motif among TE-independent Su(var)2-10 targets (data not shown), suggesting that different partners might recruit Su(var)2-10 to distinct targets. In mammals, a large family of transcription factors, the KRAB-ZFPs, are responsible for the recruitment of SetDB1 and installation of H3K9me3 mark on many different targets, primarily endogenous retroviruses (reviewed in Wolf et al., 2015). Individual members of KRAB-ZFP family influence distinct targets due to differences in DNA-binding specificities of their zinc-finger DNA-binding domains. Notably, SetDB1 recruitment through KRAB-ZFPs occurs through a SUMO-dependent mechanism (Ivanov et al., 2007). The KRAB-ZFP family is vertebrate-specific, and there are no known proteins in *D. melanogaster* that can recruit H3K9me3 activity. A preliminary search for direct Su(var)2-10 interactors using a yeast two-hybrid screen identified several proteins with putative DNA-binding domains (Ninova et al., manuscript in preparation). Thus, we propose that analogously to the KRAB-ZFP pathway in mammals, Su(var)2-10 may link DNA-binding proteins to the SetDB1 silencing machinery. Future studies are necessary to identify the proteins that guide Su(var)2-10 to target loci and to elucidate the mechanism of TE-independent recruitment of the silencing machinery.

## Materials and Methods

### Fly stocks and husbandry

shRNA-mediated knockdown experiments were performed to target the following genes of interest using the listed stocks obtained from the Bloomington Drosophila Stock Center encoding UASp-driven shRNAs: *su(var)2-10* (shSv210, BDSC #32956), *piwi* (shPiwi, BDSC #33724), *wde* (shWde, BDSC #33339), *CG30403* (BDSC #57286), and *white* (shWhite, BDSC #33623). For UASp-shSmt3, the anti-Smt3 shRNA sequence based on the TRiP line HMS01540 (Supplementary Table S1) was ligated into the pValium20 vector (Ni et al. 2011) using T4 DNA ligase from NEB, according to the manufacturer’s instructions and was integrated at the attP2 landing site by BestGene. UASp-shSetDB1 stocks were a gift from Julius Brennecke, and UASp-shSov stocks were obtained from Miklos Erdelyi (Jankovics et al., 2018). The expression of all constructs was driven by maternal alpha-tubulin67C-Gal4 (MT-Gal4, BDSC #7063), except for the shSUMO H3K9me3 ChIP-seq samples, where the BDSC #7062 stock encoding alpha-tubulin67C-Gal4 driver was used. All flies were maintained on standard medium at 24 °C and were supplemented with yeast for 2 to 3 days before ovary dissections.

### qRT-PCR and data analysis

Ovaries from 2-5 day-old females were hand dissected. Three biological replicates of 10 pairs of ovaries were used for each condition. RNA was extracted using TRIzol reagent (Invitrogen). Approximately 1 μg RNA was treated with DNase I (Invitrogen), and subsequently used as a template for cDNA synthesis using Super Script III Reverse Transcriptase (Invitrogen). SYBR Green qPCR was performed using MyTaq HS Mix (BioLine) with primers listed in Supplemental Table 1. **C**_T_ values were calculated from technical duplicates or triplicates. Relative gene expression was calculated using *Rp49/RpL13* as an endogenous reference. Data are presented as the averages of the biological replicates with error bars reflecting standard deviation.

### Heterochromatic gene definition

Heterochromatic genes were defined as genes residing on chromosome 4, chromosome U, chromosome U extra, chr2RHet, chr2LHet, chr3RHet, chr3LHet, and chrXHet, as well as the cytological borders of heterochromatin on chromosome arms 2R, 2L, 3R, 3L, and X listed in Riddle et al. 2011, dm3 assembly.

### RNA-seq and data analysis

RNA was extracted from dissected ovaries of 2-5 day old shW (control), shSu(var)2-10, and shSmt3 flies using TRIzol reagent (Invitrogen). Approximately 1 μg RNA was treated with DNase I (Invitrogen) followed by rRNA depletion using the RiboZero Gold kit (Illumina) supplemented with *D. melanogaster* rRNA antisense oligonucleotides. rRNA-depleted RNA was used as a starting material for library construction using the NEBNext Ultra Directional RNA Library Prep Kit for Illumina (New England Biolabs). Two biological replicates per condition were sequenced on Illumina HiSeq 2500 instrument at Millard and Muriel Jacobs Genetics and Genomics Laboratory at Caltech yielding 15-20 million single-end 50-bp reads. Reads were first filtered against *D. melanogaster* rRNA sequences (Stage and Eickbush, 2007) and 5S rRNA in dm3 annotated by RepeatMasker available via the UCSC Genome Browser, allowing three mismatches using bowtie v0.12.17 (Langmead et al., 2009). Less than 10% of all reads in each condition aligned to rRNA. The remaining reads were then aligned to the *D. melanogaster* genome (dm3) allowing up to two mismatches and a single mapping position bowtie v0.12.17 (-v 2, -m 1).

For differential expression analysis, read counts corresponding to the exonic regions of protein-coding genes were calculated using a custom python script. Genes with fewer than 10 reads in all samples were excluded from downstream analysis. For transposon expression, reads were aligned to the *D. melanogaster* genome (dm3) allowing 10,000 mapping positions and zero mismatches. TE annotations were obtained from RepeatMasker tables available on the UCSC browser (Karolchik et al., 2004). Read counts for individual transposon families were calculated using a custom Python script correcting for multiple mapping positions. The resulting gene and transposon count data was analyzed using the DESeq2 R package using default settings (Love et al., 2014). Significantly up- and down-regulated genes were considered to be those displaying >2-fold change between control and Su(var)2-10 GLKD libraries with an adjusted p-value <0.05. We also quantified gene expression by alignment to the reference transcriptome (dm3) followed by read count estimation by eXpress (Roberts and Pachter, 2013) and DESeq2 analysis; this yielded very similar results (data not shown). UCSC genome browser tracks were generated from the uniquely mapped read alignments using the UCSC utilities (Kent et al., 2010), using the total numbers of uniquely mapped reads to the genome per million as a normalization factor. For isoform and novel transcript analysis, reads were aligned to the *D. melanogaster* genome using hisat2 (v2.1.0), and *de novo* transcript assembly was performed using stringtie (v1.3.4d) according to the previously reported protocol (Pertea et al., 2016). Structure and expression levels of different isoforms were visualized using the ballgown R package (Frazee et al., 2015).

Euchromatic genes were classified as TE-independent if the following criteria were met: no non-reference TE insertion within 1 kb of gene ends and no more than 10% and 1000 bp of the intronic sequence and 500 bp of the 10-kb flaking regions sharing homology to TE elements as determined by RepeatMasker. To test if piRNAs mapped within gene loci, we aligned 23-29-nt reads from small RNA-seq libraries from mtGal4>UASp-shW ovaries to the genome allowing multiple mapping positions and calculated the number of reads that aligned within gene bodies and 1-kb flanking regions.

### ChIP-seq and data analysis

ChIP-seq experiments were performed as described previously (Ninova et al., accompanying manuscript) with the following antibodies: anti-H3K9me3 (Abcam, ab8898), anti-RNA Pol II (Abcam, ab5408), and anti-H3K4me2/3 (Abcam, ab6000). ChIP-seq library construction was performed using the NEBNext ChIP-Seq Library Prep Master Mix Set (NEB). Libraries were sequenced on the Illumina HiSeq 2000/2500 platform, generating single-end 49-bp or 50-bp reads. Reads were aligned to the *D. melanogaster* genome (dm3) allowing up to two mismatches and single mapping position using bowtie v0.12.17 (-v 2, -m 1). In order to determine H3K9me3-rich regions present across fly strains, we processed previously published H3K9me3 data from the ovaries of other *D. melanogaster* strains including nanos-gal4>dsWhite (GSE71374, Yu et al., 2015), nanos-gal4>dsPanx (GSE71374, Yu et al. 2015), and DGRC#204406 heterozygous strain (GSE46009, Muerdter et al., 2013).

UCSC genome browser tracks were generated as for RNA-seq tracks. H3K9me3 coverage heatmaps and average ChIP profiles on gene bodies were performed using the ngs.plot R package (Shen et al., 2014) using input libraries for normalization. For genome-wide H3K9me3 enrichment, the dm3 genome was partitioned into 5-kb intervals. Interval coverage was calculated as the ratio of RPKM-normalized reads from IP libraries to input libraries using custom python and R scripts. Low coverage intervals with fewer than 1 RPKM in input libraries were excluded from the analysis. H3K9me3-enriched 5-kb genomic intervals were defined as intervals that had H3K9me3 ChIP/Input enrichment ratio >2 in both replicate control ovaries (shW), as well as >1.5 enrichment in control ovaries of the additional analyzed datasets (shW control of shSUMO, nanos-gal4>dsWhite from GSE71374, DGRC#204406 heterozygous from GSE46009). Circular plots were generated using Circos 0.67.7 (Krzywinski et al., 2009). H3K9me3 enrichment over gene bodies was calculated as the ratio of the RPKM-normalized ChIP and input read coverage within annotated *D. melanogaster* gene ends. H3K9me3 peaks were called using macs14 using the broad peak calling protocol with the parameters (Feng et al., 2012) –nomodel, –shiftsize 73, –B, –S, and -pvalue 1e-3 using reads uniquely aligned to the dm3 *D. melanogaster* genome assembly. Peaks consistent between strains (Fig. 4A) were considered those detected independently in at least five of the seven H3K9me3 ChIP-seq data sets analyzed (four biological replicates of control UASp>shW libraries, two replicates of nanos-gal4>dsWhite from GSE71374, and one replicate of DGRC#204406 from GSE46009).

### H3K9me3 peak conservation in D. virilis

H3K9me3 ChIP-seq data from *D. virilis* ovaries was previously published (LeThomas et al., 2014). Libraries were processed using the same pipeline as for *D. melanogaster*, using the dvir-r-1.06 genome assembly version from FlyBase (St. Pierre et al., 2014). Only H3K9me3 peaks detectable in both biological replicates were considered. Orthologous genes in *D. melanogaster* and *D. virilis* were retrieved from FlyMine (Lyne et al., 2007). Genic H3K9me3 peaks of *D. melanogaster* genes were considered conserved if present within 2 kb of the annotated *D. virilis* orthologue.

### Non-reference TE insertions

Non-reference insertions were annotated using the TIDAL pipeline (Rahman et al., 2015) with default parameters using merged reads that do not map to the reference genome (allowing two mismatches) from all experiments involving DNA sequencing (DNA from Input and ChIPs) from each strain (mtGAL4>shW and mtGAL4>shSu(var)2-10).

### Co-immunoprecipitation

Expression vectors encoding GFP-fusion HP1 protein and Flag-fusion CG30403 proteins under the control of the *actin* promoter were cloned into entry cDNA clones and then transferred into pAWG or pAFW destination vectors from the *Drosophila* Gateway Vector collection using the Gateway system. Vectors were co-transfected into S2 cells, cultured in Schneider’s *Drosophila* Medium containing 10% heat-inactivated FBS and 1X penicillin-streptomycin with the TransIT-LT 1 reagent (Mirus). Flag-CG30403 only (no bait) transfected cells were used as negative control. At 24-48 hours post transfection cells were harvested and lysed in lysis buffer (20 mM Tris-HCl, pH 7.4, 150 mM NaCl, 0.2% NP-40, 0.2% Triton-X, 5% glycerol) supplemented with protease inhibitor cocktail (Roche). Lysates were incubated with GFP-Trap magnetic agarose beads (Chromotek) for 1-2 hours at 4 °C with end-to-end rotation. After incubation, the beads were washed five times with 500 μl wash buffer (0.1% NP40, 20 mM Tris, pH 7.4, 150 mM NaCl) containing protease inhibitor. Proteins were detected by Western blot using HRP-conjugated mouse anti-FLAG (Sigma, A8592) and rabbit polyclonal anti-GFP (Chen et al., 2016), followed by IRDye anti-rabbit secondary antibody (Li-cor 925-32211).

## Data availability

All RNA-seq and ChIP-seq data generated for this study and for the accompanying manuscript (Ninova et al, accompanying manuscript) were deposited into the Gene Expression Omnibus (GEO) database under accession GSE115277. H3K9me3 ChIP-seq data for shSov and respective controls was deposited to GEO under accession GSE125055.

## Acknowledgements

We thank members of the Fejes Toth and Aravin labs for discussion. We are grateful to the Bloomington Stock Center and Julius Brennecke for providing fly stocks. We thank Igor Antoshechkin (Caltech) for help with sequencing. This work was supported by grants from the National Institutes of Health (R01 GM097363), the Ministry of Education and Science of Russian Federation (14.W03.31.0007) and by the Packard Fellowship Awards to A.A.A., and the National Institutes of Health (R01GM110217) and Ellison Medical Foundation Awards to K.F.T..

## Author contributions

MN, AAA and KFT designed the experiments. MN, BG, YL, YAC and SP executed the experiments, FJ and ME generated reagents. MN performed the computational analysis and interpretation of the data. The manuscript was written by MN, AAA and KFT.

## Supplementary Materials

**Figure S1. Loss of H3K9me3 islands at euchromatic genes leads to ectopic gene expression.** Regulation of testis-biased endo-siRNA locus *CG44774/CG4068/esi-2* through a Su(var)2-10-dependent H3K9me3 mark. UCSC browser tracks show the RNA-seq and H3K9me3 ChIP-seq signals in the indicated conditions. The *CG44774* gene is marked by a Su(var)2-10- and SUMO-dependent, but Piwi-independent, H3K9me3 peak and is expressed at low levels in the ovary. Su(var)2-10 GLKD leads to H3K9me3 loss and transcript up-regulation. Areas shaded in grey represent repetitive regions where reads did not align uniquely.

**Figure S2. Sov regulates genome-wide H3K9me3 profiles.** (A) Circle plot of the H3K9me3 genome-wide distribution and fold changes upon Sov GLKD. Outer circle shows the enrichment of H3K9me3 (IP/Input) for 5-kb genomic windows in control ovaries. Black tiles indicate 5-kb windows enriched in H3K9me3 (>2 IP/Input signal ratio). Inner circle shows the fold change of H3K9me3 signal between Sov KD and control ovaries for H3K9me3-enriched regions. (B) Boxplot of the log2-transformed fold change of H3K9me3 signal upon Sov GLKD relative to control ovaries for 5-kb genomic windows enriched or depleted of H3K9me3 signal (>2 fold enrichment cutoff). (C) CG30403 interacts with HP1. Flag-CG30403 and GFP-HP1a were co-expressed in S2 cells. Western blot shows results from co-immunoprecipitation using GFP-HP1 as a bait.

**Table S1. List of primers used in RT-qPCR experiments.**

