## Supplemental_figures_and_table for "The SUMO ligase Su(var)2-10 controls eu- and heterochromatic gene expression via establishment of H3K9 trimethylation and negative feedback regulation"

### Ninova et al., Figure S1

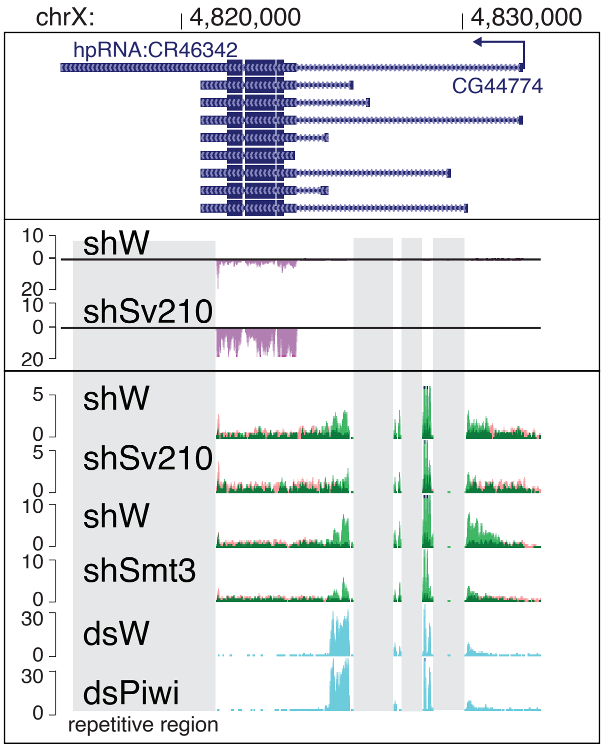

Ninova et al., Figure S2

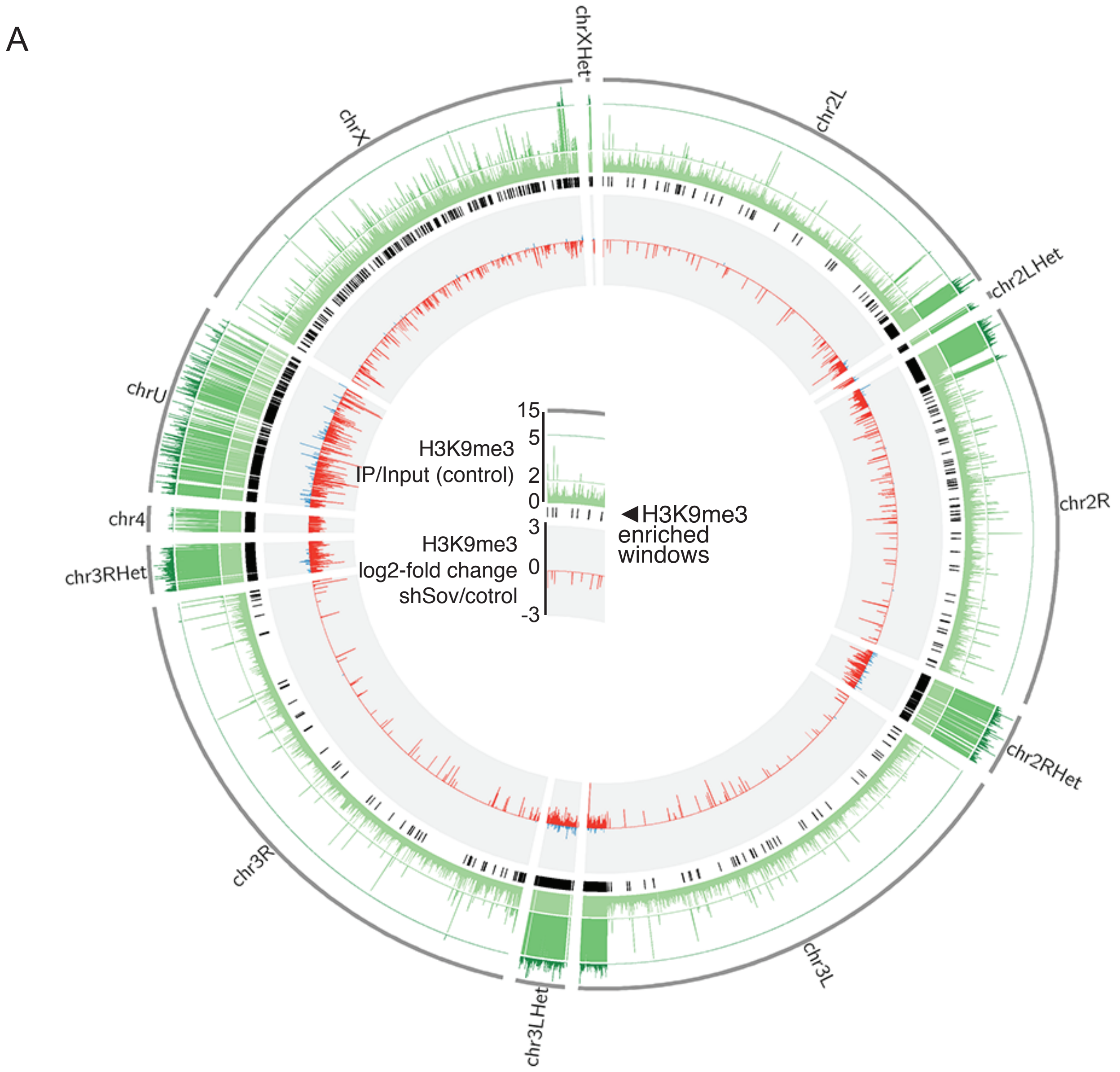


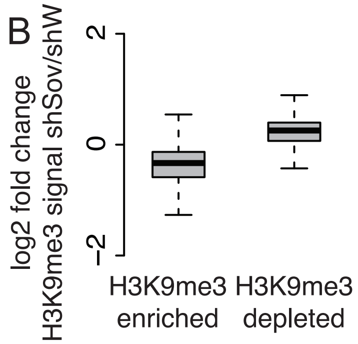


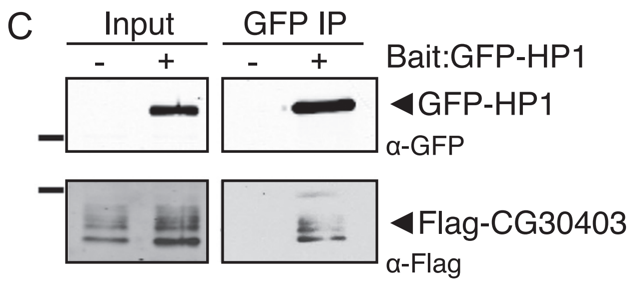

Supplementary Table 1. List of primers used in RT-qPCR

| | | |
|---|---|---|
| Wde (long isoform) | AAACGAGTTTGTCGTGTC | TGGTCTGGTTTACTCCCATCA |
| Wde (nascent transcript) | GAGCAGGGATCCTCATTGAA | GCTGCCTGTTTCCTCATCTC |
| Sov (all isoforms) | AATGTTACCGCCAATGGAAG | TTCCCGTTTTGATTGCTAGG |
| Sov (long isoform) | CCTTATGGCCAGCTCCTTCG | TTGCCGGCGAATACGATAGA |
| Sov (nascent transcript) | GGAATGTGCATCAACGACGA | AGTCGCAAAAGCTCGTCCAC |
| Smt3 | ACCATCGAGGTTTACCAGCA | TGTGTTTTTGCTTTTGTGGTTT |
| CG30403 | TGGTTTGCCCCACTTCACG | GGCGCTTTAGTTGAGGAATTGAT |
